# MIAMI: Mutual Information-based Analysis of Multiplex Imaging data

**DOI:** 10.1101/2022.02.10.479967

**Authors:** Souvik Seal, Debashis Ghosh

## Abstract

**Motivation:** Studying the interaction or co-expression of the proteins or markers in the tumor microenvironment (TME) of cancer subjects can be crucial in the assessment of risks, such as death or recurrence. In the conventional approach, the cells need to be declared positive or negative for a marker based on its intensity. For multiple markers, manual thresholds are required for each marker, which can become cumbersome. The performance of the subsequent analysis relies heavily on this step and thus suffers from subjectivity and lacks robustness.

**Results:** We present a new method where different marker intensities are viewed as dependent random variables, and the mutual information (MI) between them is considered to be a metric of co-expression. Estimation of the joint density, as required in the traditional form of MI, becomes increasingly challenging as the number of markers increases. We consider an alternative formulation of MI which is conceptually similar but has an efficient estimation technique for which we develop a new generalization. With the proposed method, we analyzed a lung cancer dataset finding the co-expression of the markers, HLA-DR and CK to be associated with survival. We also analyzed a triple negative breast cancer dataset finding the co-expression of the immuno-regulatory proteins, PD1, PD-L1, Lag3 and IDO, to be associated with disease recurrence. We demonstrated the robustness of our method through different simulation studies.

**Availability:** The associated *R* package can be found here, https://github.com/sealx017/MIAMI.

**Supplementary information:** The Supplementary Material is attached.

## 1 Introduction

In recent years, multiplex tissue imaging (Bataille *et al*., 2006) technologies like, imaging mass cytometry (IMC) (Ali *et al*., 2020), multiplex immunohistochemistry (mIHC) (Tan *et al*., 2020) and multiplexed ion beam imaging (MIBI) (Angelo *et al*., 2014) have become increasingly popular for probing single-cell spatial biology. The technologies help in understanding the biological mechanisms underlying cellular and protein interactions in a wide array of scientific contexts. MIBI (Ionpath Inc.) (Angelo *et al*., 2014) platform and the mIHC platforms such as Vectra 3.0 (Akoya Biosciences) (Huang *et al*., 2013) and Vectra Polaris (Akoya Biosciences) (Pollan *et al*., 2020) produce images of similar structure. In particular, each image is two-dimensional, collected at cell- and nucleus-level resolution and proteins in the sample have been labeled with antibodies that attach to cell membranes. We will refer to the antibodies as markers in the paper. Typically, mIHC images have 6-8 markers, whereas MIBI images can have 40 or more markers.

Many of the above markers are surface or phenotypic markers (Zola *et al*., 2007; Shipkova and Wieland, 2012) which are primarily used for cell type identification. Additionally, there are functional markers (Ijsselsteijn *et al*., 2019) such as HLA-DR (Saraiva *et al*., 2018), PD1, PD-L1 (Alsaab *et al*., 2017) and CD45RO (Lee *et al*., 2008) that dictate or regulate important cell-functions. Both surface and functional markers are quantified as continuous valued marker intensities. However, in the traditional method, the interaction or co-expression effects of the markers are studied by binarizing them. For every marker, a threshold is chosen to indicate whether a cell (or, a pixel) in the tumor microenvironment (TME) (Binnewies *et al*., 2018) is positive or negative for that marker. If a cell is positive for two markers, it implies that the markers have co-expressed or co-occurred in that cell. Next, for all the subjects considering every unique pair of markers, the proportion of cells positive for both the markers, the proportion of cells positive for only the first marker and the proportion of cells positive for only the second marker are computed. These proportions can then be used in hiearchical clustering (Murtagh and Legendre, 2014) to group the subjects. It can then be tested if the cluster labels correlate with clinical outcomes, such as disease recurrence or time to death (Koguchi *et al*., 2015; Johnson *et al*., 2021; Jackson *et al*., 2020). The step of binarizing the marker expression profiles can be performed either using compatible commercially available software or manual assessment of the segmented images. For example, Johnson *et al*. (2018) used AQUAnalysis® software (McCabe *et al*., 2005; Dolled-Filhart *et al*., 2010) for binarizing the functional markers, PD1, PDL1, HLA-DR and IDO, and discovered that their co-expression predicted improved outcomes of anti-PD-1 therapies in metastatic melanoma. On the other hand, Patwa *et al*. (2021) used their own clinical expertise and careful evaluation of the segmented images to determine the binarizing thresholds and discovered the interaction between the markers, PD-1, PD-L1, IDO, and Lag3 to be associated with recurrence. It should be pointed out that such expression-thresholding or binarization is also popular in cell-cell communication or interaction analysis (Jin *et al*., 2021) in the context of single-cell RNA sequencing (scRNA-seq) data. As an example, Armingol *et al*. (2021) defined a pair of cells to be interacting if the expressions of both ligand and receptor (Wong *et al*., 1997) in those cells exceed certain chosen thresholds.

The manual threshold selection process for multiple markers can be extremely challenging and is subjective. Alternatively, commercially available softwares, such as AQUAnalysis® and inForm (Dolled-Filhart *et al*., 2010; Kramer *et al*., 2018), can be used for automated threshold selection in the context of a few particular data types. However, these softwares generally follow a “black box approach” and can be difficult to interpret. On top of these difficulties, many authors have criticized binarizing continuous random variables in general due to the resulting loss of power (Irwin and McClelland, 2003; Altman and Royston, 2006).

In this paper, we propose a threshold-free approach for studying marker co-expression. We treat the marker intensities as continuous random variables and use their marginal and joint probability density functions (PDF’s) to construct a metric of co-expression based on mutual information (MI) (Cover and Thomas, 2006). Unlike the correlation coefficient, MI is capable of capturing non-linear patterns of dependence between the markers and is easily extendable for more than two markers. However, as the number of markers increases, computing the joint PDF becomes increasingly challenging which makes the computation of MI infeasible as well. Therefore, we use a slightly different formulation of MI known as Euclidean quadratic mutual information (EQMI) (Principe *et al*., 2000) which has a similar interpretation but can be computed more efficiently. The computation algorithm is discussed in Principe (2010) with a simpler assumption that we further generalize. The vector of estimated values of the EQMI of all the subjects is tested for association with clinical outcomes. With the proposed method, we analyzed an mIHC lung cancer dataset (Seal *et al*., 2022) finding that a higher co-expression of the markers, HLA-DR and CK was significantly associated with better five year overall survival. We analyzed a MIBI triple-negative breast cancer (TNBC) dataset (Keren *et al*., 2018) studying the co-expression of two sets of functional markers, (a) HLA-DR, CD45RO, H3K27me3, H3K9ac and HLA-Class-1, and (b) PD1, PD-L1, Lag3 and IDO, which are also known as immuno-regulatory proteins (IRP’s). We found the co-expression of the IRP’s to be significantly associated with recurrence. We demonstrated the robustness of our method over the existing approaches through different simulation studies.

## 2 Materials and Methods

Suppose there are *p* markers and *N* subjects with the *j*-th subject having *n*_*j*_ cells. Let *X*_*kij*_ denote the expression of the *k*-th marker in the *i*-th cell of *j*-th subject for *k* = 1, 2, …, *p, i* = 1, 2, …, *n*_*j*_, and *j* = 1, 2, …, *N*. Note that we focus on cell-level data in this paper but the framework is readily usable on pixel-level data as well. Let *Y* = (*Y*_1_, *Y*_2_, …, *Y*_*N*_)^*T*^ denote a subject-level outcome vector and *C* be an *N* ×*S* matrix of subject-level covariates. Next, we discuss the existing and proposed methods and a brief summary of both is provided in Figure 1.

**Fig. 1.**
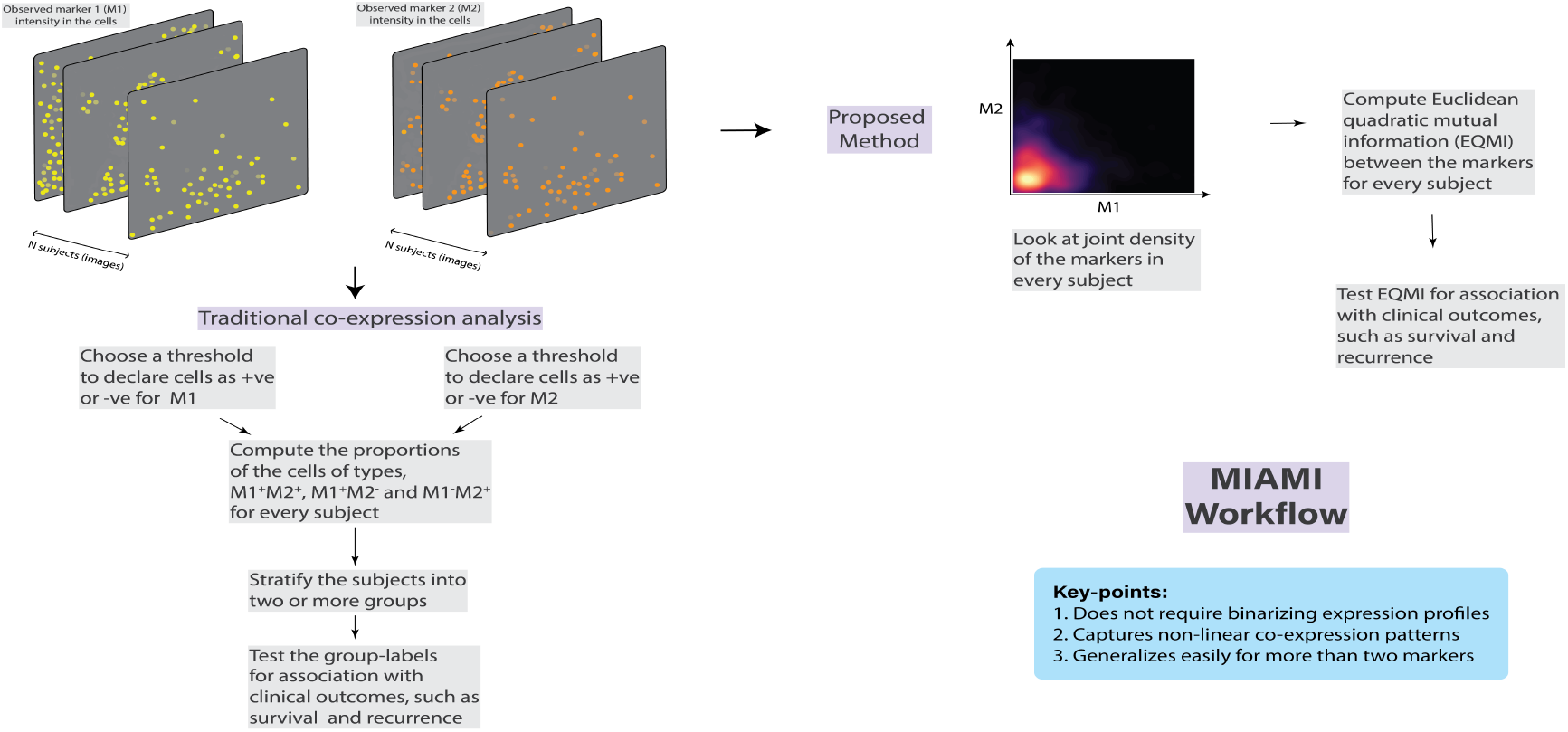
Comparison of the workflow of the proposed method with the traditional method. We used segmented cell-level data in this paper but the method is applicable on a pixel-level data as well.

### 2.1 Traditional thresholding-based approach to study co-expression of the markers

For every marker *k*, we choose a cut-off *t*_*k*_ to define cell *i* of subject *j* as positive for that marker if *X*_*kij*_ *> t*_*k*_. The choice of *t*_*k*_ can be guided by prior biological insight or by careful inspection of the marker intensity profile. For example, extreme quantiles like (e.g., the 90th or 95th percentiles) can serve as viable thresholds, as we see in the simulations. However, in most real datasets, it requires user-defined thresholds to obtain an appropriate threshold that leads to meaningful and interpretable conclusion. A cell can be positive for multiple markers. If a cell is positive for a pair of markers, (*k*_*r*_, *k*_*s*_), it would imply that these markers have co-expressed in that particular cell. For every subject, compute the proportion of such double-positive cells, denoted by 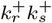, the proportion of cells positive for only the first marker, denoted by 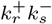, and the proportion of cells positive for only the second marker, denoted by 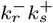, for *k*_*r*_, *k*_*s*_ ∈ {1, 2, …, *p*} and *k*_*r*_ ≠ *k*_*s*_. Next, the subjects are classified into two or more groups based on their vectors of proportions using hierarchical clustering. Note that if it is biologically relevant, instead of a pairwise analysis, one can also study multiple markers jointly e.g., with four markers, one can count the cells which are either 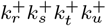 or any of the possible (2^4^ − 1) = 15 combinations and group the subjects based on these proportions.

Suppose that the subjects are grouped into *M* clusters. Let **Z** = (*Z*_1_, …, *Z*_*N*_)^*T*^ be an *N* × *M* matrix of the cluster labels. *Z*_*j*_ is a vector corresponding to the subject *j* with *Z*_*jm*_ = 1 if the *j*-th subject belongs to group *m* and 0 otherwise. In most common practices, *M* = 2 is considered. When *Y* is a continuous outcome, a standard multiple linear regression model with **Z** as a predictor can be written as

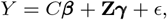

where ***β, γ*** are fixed effects and *ϵ* is an *N* × 1 error vector following multivariate normal distribution (MVN) with mean **0** and identity covariance matrix *σ*^2^𝕀_*N*_. The null hypothesis, *H*_0_ : ***γ*** = **0**, can be tested using the Wald test or likelihood ratio test (LRT) (Gourieroux *et al*., 1982). Similarly, when *Y* is a categorical outcome, a logistic or multinomial logistic regression model (Kwak and Clayton-Matthews, 2002) can be considered.

Next, we consider the case of *Y* being a a right-censored failure time outcome. Let the outcome of the *j*-th individual be *Y*_*j*_ = *min*(*T*_*j*_, *U*_*j*_), where *T*_*j*_ is the time to event and *U*_*j*_ is the censoring time. Let *δ*_*j*_ ≡ *I*(*T*_*j*_ ≤ *U*_*j*_) be the corresponding censoring indicator. Assuming that *T*_*j*_ and *U*_*j*_ are conditionally independent given the covariates for *j* = 1, 2, …, *N*, the hazard function for the Cox proportional hazards (PH) model (Andersen and Gill, 1982) with fixed effects can be written as,

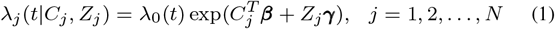

where *λ*_*j*_ (*t*|*C*_*j*_, *Z*_*j*_) is the hazard of the *j*-th subject at time *t*, given the vector of covariates *C*_*j*_ and the cluster label *Z*_*j*_ and *λ*_0_(*t*) is an unspecified baseline hazard at time *t*. To test the null hypothesis: *H*_0_ : ***γ*** = **0**, an LRT (Therneau, 1997) can be considered.

### 2.2 Proposed Method: Mutual Information based analysis of marker co-expression

#### 2.2.1 Theory of Mutual Information

Mutual information (MI) is an information theoretic measure of dependence between two or more random variables (r.v.’s). In contrast to the linear correlation coefficient, it captures dependences which do not manifest themselves in the covariance (Kraskov *et al*., 2004). For two random variables, *R*_1_ and *R*_2_ with sample space 𝒟, marginal probability density functions (PDFs), *f*_1_, *f*_2_, and joint PDF *f*_12_, the MI is defined as the following:

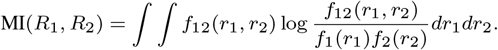

MI has been used as a tool for feature selection in many different contexts (Yang and Moody, 1999; Liu *et al*., 2009; Hoque *et al*., 2014; Song *et al*., 2021). The measure can be easily generalized for more than two random variables. However, due to the curse of dimensionality (Langrené and Warin, 2019), it becomes extremely difficult to estimate the joint PDF and hence, the MI, as the number of random variables increases.

Xu (1998) and Principe *et al*. (2000) looked into alternative definitions of MI that would remain conceptually similar but can be computed efficiently. They discussed two measures, Euclidean quadratic mutual information (EQMI) and Cauchy-Schwarz quadratic mutual information (CSQMI), defined as below,

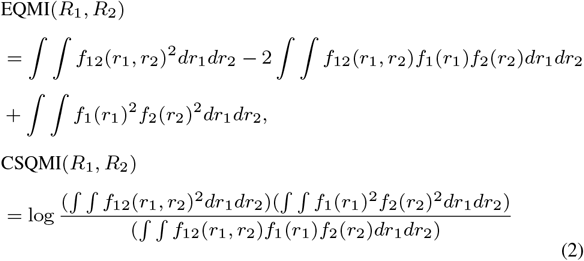

It is trivial to verify that EQMI(*R*_1_, *R*_2_) ≥ 0 with equality occurring if and only if *R*_1_, *R*_2_ are independent i.e., *f*_12_(*r*_1_, *r*_2_) = *f*_1_(*r*_1_)*f*_2_(*r*_2_) for any *r*_1_, *r*_2_ ∈ ℝ. Using the Cauchy-Schwarz inequality, we have CSQMI(*R*_1_, *R*_2_) ≥ 0 with equality happening if and only if *R*_1_, *R*_2_ are independent. Xu (1998) argued that EQMI shares more properties, such as convexity with respect to the PDF’s, of the traditional MI compared to CSQMI and thus, is better suited as a dependence measure. It should also be noticed that the form of EQMI is very similar to the distance covariance measure proposed by Székely *et al*. (2007). There is one more generalized measure of dependence, known as kernel canonical correlation analysis (KCCA) (Huang et al., 2006), which does not share forumlaic similarity with EQMI but can potentially serve as an alternative for detecting non-linear dependence patterns. In this paper, we focus on EQMI and propose its usage in the co-expression analysis of the markers in general multiplex imaging datasets used in the study of spatial biology of TME.

#### 2.2.2 Formulation of EQMI

For every subject *j*, we assume that the expression of marker *k* is a continuous random variable, denoted by *X*_*kj*_, with sample space 𝒟 = [0, 1]. *X*_*kj*_ is observed in *n*_*j*_ cells as, 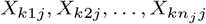. Suppose there are *p* = 2 markers and for every subject *j*, denote their joint PDF as *f*_12*j*_ (*x*_1_, *x*_2_) and their marginal PDFs as *f*_1*j*_ (*x*_1_) and *f*_2*j*_ (*x*_2_) respectively. Following Equation (2), the EQMI between the markers 1 and 2 for subject *j* can be defined as,

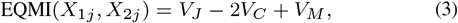

where *V*_*J*_ = ∫ ∫ *f* _12*j*_ (*x*_1_, *x*_2_)^2^*dx*_1_ *dx*_2_, *V*_*C*_ = ∫ ∫ *f* _12*j*_ (*x*_1_, *x*_2_)*f*_1*j*_ (*x*_1_)*f* (*x*)*dx dx*, and *V*_*M*_ = ∫ ∫ *f*_1*j*_ (*x*_1_)^2^*f*_2*j*_ (*x*_2_)^2^*dx*_1_*dx*_2_. EQMI(*X*_1*j*_, *X*_2*j*_) can be interpreted as a generalized measure of co-expression of the markers, capable of capturing non-linear dependences. A large value of EQMI(*X*_1*j*_, *X*_2*j*_) will imply that the markers 1, 2 have significantly co-expressed in the TME of subject *j*. EQMI(*X*_1*j*_, *X*_2*j*_) is bounded below by 0 for every *j* but there is no common upper bound for different *j*’s. Therefore, to compare EQMI(*X*_1*j*_, *X*_2*j*_)’s across different subjects, we need to appropriately standardize the values so that they lie in the same scale. We define a new measure as, EQMI*\**(*X*_1*j*_, *X*_2*j*_) = (*V*_*J*_ + *V*_*M*_)^−1^(*V*_*J*_ − 2*V*_*C*_ + *V*_*M*_). We observe that EQMI*\**(*X*_1*j*_, *X*_2*j*_) lies between [0, 1] because *V*_*C*_ ≥ 0, and has a similar interpretation as EQMI(*X*_1*j*_, *X*_2*j*_).

It is intuitive to generalize the measure for any *p* ≥ 2 markers. For subject *j*, letting *f*_12…*pj*_ (*x*_1_, *x*_2_, …, *x*_*p*_) to be the joint PDF and *f*_1*j*_ (*x*_*p*_), *f*_2*j*_ (*x*_*p*_), …, *f*_*pj*_ (*x*_*p*_) to be the marginal PDFs, EQMI*\** can be defined as,

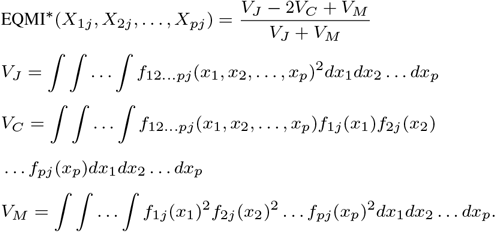

Notice that to estimate EQMI*\**(*X*_1*j*_, *X*_2*j*_, …, *X*_*pj*_) when the true PDFs are unknown, a naive approach would be to estimate the PDFs first by using a kernel density estimation (KDE) approach (Silverman, 1981). Then, we use the estimated PDFs to compute the terms *V*_*J*_, *V*_*C*_ and *V*_*M*_ via numerical integration (Davis and Rabinowitz, 2007). However, such an approach would be computationally infeasible for a large *p* and will defeat the purpose of considering EQMI instead of the standard form of MI.

EQMI*\**(*X*_1*j*_, *X*_2*j*_, …, *X*_*pj*_) can be estimated efficiently as, 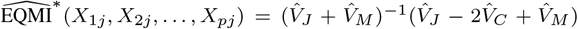, where

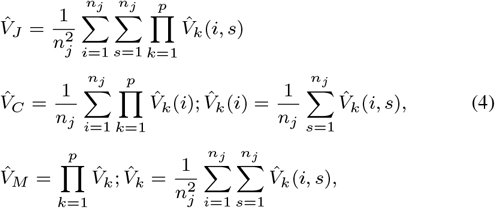

and 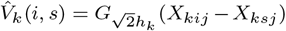. *G*_*h*_ stands for a Gaussian kernel with bandwidth parameter *h*. This clever way of estimating the terms, 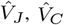 and 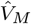 is described in Principe (2010) with a simpler assumption that *h*_*k*_’s are equal for all *k* i.e., *h*_*k*_ = *h* for *k* = 1, 2, …, *p*. We provide the general derivation (i.e., *h*_*k*_ ≠ *h*_*k′*_, for *k ≠ k′*) and other associated details in the Supplemetary material. For choosing the optimal values of *h*_*k*_’s, we generally consider the diagonal multivariate plug-in bandwidth selection procedure (Wand *et al*., 1994; Chacón and Duong, 2010). However, when *p* is large (*p >* 6), to avoid computational deadlock we suggest using Silverman’s rule of thumb (Silverman, 1981) for choosing *h*_*k*_’s individually.

#### 2.2.3 Using Mutual Information in association analysis

Denote the vector of estimated values of EQMI*\** as **E** ≡ (*E*_1_, …, *E*_*N*_)^*T*^, where 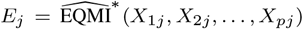. Our goal is to test if **E** is associated with the clinical outcome *Y*. When *Y* is a continuous outcome, a standard multiple linear regression model with **E** as a predictor can be written as,

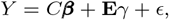

where ***β***, *γ* are fixed effects and *ϵ* is an *N* × 1 error vector following multivariate normal distribution (MVN) with mean **0** and identity covariance matrix *σ*^2^𝕀_*N*_. After estimating the parameters, the null hypothesis, *H*_0_ : *γ* = 0, can be tested using the Wald test. Note that we can also add higher order terms of *E*_*j*_, such as 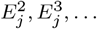 as in a polynomial regression model (Ostertagová, 2012) to account for non-linear relationship between *Y* and *E*_*j*_. Similarly, when *Y* is a categorical outcome, a logistic or multinomial logistic regression model (Kwak and Clayton-Matthews, 2002) can be considered.

Next, we consider the case of *Y* being a survival or recurrence outcome. Using the same definitions and conditional independence assumptions of *T*_*j*_, *U*_*j*_ and covariates as in Section 2.1, the hazard function for the Cox proportional hazards (PH) model can be written as,

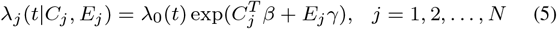

where *λ*_*j*_ (*t*|*C*_*j*_, *E*_*j*_) is the hazard of the *j*-th subject at time *t*, given the vector of covariates *C*_*j*_ and the predictor *E*_*j*_ and *λ*_0_(*t*) is an unspecified baseline hazard at time *t*. To test the null hypothesis: *H*_0_ : *γ* = 0, a likelihood ratio test (LRT) can be considered.

In a two-marker scenario (i.e., *p* = 2), one can also treat the absolute value of the Pearson correlation as a measure of marker co-expression and use it as *E*_*j*_ in the earlier equations for testing association with the outcome. However, as we later demonstrate, using correlation instead of EQMI can be sub-optimal in many cases. It is also difficult to generalize to more than two markers.

## 3 Real Data Analysis

We applied our method on two real datasets, an mIHC lung cancer dataset (Seal *et al*., 2021) from Vectra 3.0 platform and a MIBI triple-negative breast cancer dataset (Keren *et al*., 2018). We also applied the traditional thresholding-based method on both the datasets. Since it was hard to decide the optimal thresholds for binarizing the markers, we ran the method for varying values of the thresholds. For every marker, concatenating the intensity data of all the subjects, we computed the median, 95% and 99% quantiles. Next, three different thresholding-based methods using these quantiles were considered, respectively referred to as Median-Thresholding, Threshold 1 and Threshold 2. Note that Threshold 1 and 2 both captured the difference in the tails of the distributions, whereas Median-Thresholding captured the difference in the centers. For the first dataset, we also performed the correlation-based association analysis, referred to as Corr.

### 3.1 Application to mIHC Lung Cancer data

In the lung cancer dataset, there were 153 subjects each with 3-5 images (in total, 761 images). Two subject-level covariates, age and sex were available. For every subject, the images were non-overlapping and from the same tumor microenvironment (TME) region. After segmenting the images, the subjects had varying number of identified cells (from 3,755 to 16,949). We worked directly with the cell-level data as described next. The cells come from two different tissue regions: tumor and stroma. They were classified into either of the six different cell types: CD14+, CD19+, CD4+, CD8+, CK+ and Other, based on the expression of the phenotypic markers, CD19, CD3, CK, CD8 and CD14. These markers (and the cell-types) usually have clinical meaning e.g, CK is a type of a tumor cell marker and CD4 and CD8 are T-cell or cytotoxic T-cell markers. Apart from these, a functional marker HLA-DR (also known as MHCII), was measured in each of the cells. Johnson *et al*. (2021) classified the subjects into two groups, (a) MHCII: High and (b) MHCII: Low based on the proportion of CK^+^ tumor cells that are also positive for HLA-DR (i.e., CK^+^HLA-DR^+^ cells). They discovered that group (a) had significantly higher five-year overall rate of survival (reported *p*-value of 0.046).

Note that having a large number of CK+HLADR+ tumor cells implies that these two markers had co-expressed in a lot of the tumor cells. In light of that, we studied if the degree of co-expression of these markers in the tumor cells, as quantified by our method, was associated with the survival. Considering the tumor cells of every subject *j*, we first estimated EQMI*\**, 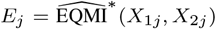 between the two markers, HLA-DR and CK. Next, we tested the association of *E*_*j*_ with five-year overall survival using the Cox-PH model from Equation (5). The coefficient *γ* was -7.26 with the *p*-value of the LRT being 0.0286. Thus, subjects with high co-expression of the markers in the tumor cells were more likely to survive. The result was thus consistent with Johnson *et al*. (2021)’s finding. The estimated coefficient, hazard ratio (HR) and *p*-value of all the methods are listed in Table 1 and further details such as confidence interval of the HR, are provided in the Supplementary Material. The correlation-based analysis (Corr) yielded a negative coefficient estimate but was not statistically significant (at level 0.05). Out of the thresholding-based methods, only Threshold 2 had a significant *p*-value. It demonstrated that the traditional thresholding-based method could vary heavily based on the choice of the thresholds leading to utterly different conclusions. On the other hand, inference using our method would be robust.

**Table 1.** Estimated coefficient, hazard ratio (HR) and LRT *p*-value for testing association with five-year overall survival using different methods in the mIHC lung cancer dataset.

| Method | Coefficient (HR) | LRT $p$-value |
| --- | --- | --- |
| EQMI* | -7.26 (0.0007) | 0.0286 |
| Corr | -1.38 (0.2512) | 0.1838 |
| Median-Thresholding | -0.48 (0.6189) | 0.0495 |
| Threshold 1 | -0.58 (0.5585) | 0.0575 |
| Threshold 2 | -1.03 (0.3555) | 0.0098 |

We should point out that CK was a phenotypic marker, and studying its co-expression with a functional marker HLA-DR might not have much clinical relevance. However, the goal was to demonstrate that our method obviates the need to binarize the continuous-valued marker expression profiles and is potentially applicable even when one of the markers is a phenotypic marker.

### 3.2 Application to TNBC MIBI data

The triple-negative breast cancer (TNBC) MIBI dataset (Keren *et al*., 2018) had 38 subjects, each with one image. There were 201,656 cells in total with each of the images having varying numbers of cells (between 1217 and 8212). There were 49 markers in total, the majority of which were lineage or phenotypic markers, used primarily for cell-type identification. We were interested in two sets of functional markers, (a) HLA-DR, CD45RO, H3K27me3, H3K9ac and HLA-Class-1, and (b) PD1, PD-L1, Lag3 and IDO, also known as immuno-regulatory proteins (IRP’s). The reason for concentrating on these two sets of markers was the findings of Patwa *et al*. (2021). Employing the thresholding-based method with a set of very carefully chosen thresholds, they concluded that the pair-wise co-expression of the functional markers from the sets (a) and (b) were negatively associated with two clinical outcomes, namely disease recurrence and time to death (survival). For set (a), they did not find any statistical significance for either of the clinical outcomes (at level 0.05), whereas, for set (b), they were able to find statistical significance in the association test with recurrence (reported *p*-value of 0.0058).

For the co-expression analysis with the five markers from the set (a), we looked into all possible (2^5^ − 5 − 1 = 26) two-way and higher-order combinations of the markers. There were four different types of combination, namely pair (10), triplet (10), quadruplet (5) and quintuple (1). We computed EQMI*\** for each combination of the markers and tested for association with recurrence and survival. In Table 2, we list the results for five marker combinations for which the lowest *p*-values were observed. We noticed that at a level of 0.05, several marker-combinations were found to be associated with recurrence. All the estimated coefficients were negative, implying that higher co-expression of the markers decreased the chance of disease recurrence. However, once we adjusted the *p*-values for multiple testing correction using Bonferroni’s method (Bonferroni, 1936), only the combinations “HLA-DR, CD45RO” and “HLA-DR, CD45RO, H3K9ac” remained significant. Note that, we compared the *p*-values of every type of combination separately, meaning that for marker pairs and triplets, we compared the *p*-values at level 0.05/10 since the numbers of pairs and triplets were both 10, and for quadruplets, at level 0.05/5 since the number of quadruplets was 5. The marker-combinations were not independent and thus, a Bonferroni correction probably has been overly conservative in this case. For the survival outcome, we did not detect any statistical significance. But, the negative coefficient estimates hinted at a possible association of better rate of survival with higher co-expression.

**Table 2.** Estimated coefficient and LRT *p*-value for testing association with recurrence and survival for five combinations of the markers from sets (a) and (b) with the lowest *p*-values, obtained by the proposed method in the MIBI dataset.

| Clinical Outcome | Marker Combination | Coefficient | LRT $p$-value |
| --- | --- | --- | --- |
| Recurrence | HLA-DR, CD45RO | -32.97 | 0.0022 |
|  | HLA-DR, CD45RO, H3K9ac | -12.48 | 0.0051 |
|  | HLA-DR, CD45RO, H3K27me3 | -6.73 | 0.0245 |
|  | CD45RO, H3K9ac | -41.53 | 0.0329 |
|  | HLA-DR, CD45RO, HLA-Class-1 | -7.20 | 0.0421 |
| Survival | HLA-Class-1, H3K27me3 | -12.75 | 0.1010 |
|  | HLA-DR, H3K9ac | -8.27 | 0.1542 |
|  | HLA-DR, CD45RO | -8.35 | 0.1583 |
|  | HLA-DR, HLA-Class-1 | -29.69 | 0.1630 |
|  | HLA-DR, CD45RO, H3K9ac | -0.74 | 0.3158 |
| Recurrence | PD1, PD-L1 | -9.61e+03 | 0.0046 |
|  | PD1, PD-L1, IDO | -5.54e+02 | 0.0065 |
|  | PD1, PD-L1, Lag3, IDO | -4.37e+02 | 0.0069 |
|  | PD-L1, Lag3, IDO | -8.94e+02 | 0.0084 |
|  | PD1, PD-L1, Lag3 | -1.91e+02 | 0.0090 |
| Survival | PD-L1, Lag3 | -1.20e+02 | 0.0103 |
|  | Lag3, IDO | -1.14e+02 | 0.0302 |
|  | PD-L1, Lag3, IDO | -5.67e+02 | 0.0449 |
|  | PD-L1, IDO | -6.85e+02 | 0.0490 |
|  | PD1, PD-L1, Lag3, IDO | -2.15e+02 | 0.0586 |

For the co-expression analysis with the four IRP’s from the set (b), we looked into all possible (2^4^ − 4 − 1 = 11) two-way and higher order combinations. There were four different types of combination, namely pair (6), triplet (4) and quadruplet (1). In Table 2, we list the results for five marker combinations for which the lowest *p*-values were observed. At a level of 0.05, many of the marker-combinations were found to be associated with both recurrence and survival. Again, all the estimated coefficients were negative, implying that higher co-expression of the markers decreased the chance of disease recurrence. Upon correcting the *p*-values for multiple testing using Bonferroni’s method, all of the marker combinations listed in Table 2 remained significant for recurrence while for survival, none of them remained significant. We compared the *p*-values of every type of combination separately, meaning that for marker pairs we compared the *p*-values at level 0.05/6 since the number of pairs was 6, for triplets we compared the *p*-values at level 0.05/4 since the number of triplets was 4, and for quadruplet, at level 0.05 since there was just a single quadruplet. Thus, we also arrived at a similar conclusion as (Patwa *et al*., 2021) that the inter-play or co-expression of the IRP’s were significantly associated with recurrence and possibly also with survival. One added novelty of our method was that one could easily pinpoint which of the combinations of the IRP’s had the most impact.

It should be kept in mind that the sample-size for this dataset was quite small with only 16 events for recurrence and 15 for survival. It might have affected the overall inference which was based on asymptotic distributional properties of the test statistics. For the same reason, we mainly focused on the sign of the coefficent estimates but not their CI’s in this particular case. We also applied the simple thresholding-based methods, Median-Thresholding, Threshold 1 and Threshold 2 as described earlier using both the sets of markers. We found only a single statistically significant result which was for Median-Thresholding using set (b) in the association test with recurrence. Associated tables are provided in the Supplementary Material.

## 4 Simulation

Next, we compared the performance of the EQMI*\**-based association analysis with the correlation-based association analysis and the thresholding-based methods in different simulation setups. We assumed that there were two groups of subjects, one with subjects having high marker co-expression and the other with subjects having low or almost zero marker co-expression. In Section 4.1, we considered two markers sharing linear and non-linear patterns of co-expression relationship. In Section 4.2, we considered three markers sharing varying degree of linear co-expression relationship.

### 4.1 Simulation with two markers

#### 4.1.1 Simulation using Gaussian copula

We replicated the characteristics of the lung cancer dataset in this simulation. The mean marginal distribution of the markers HLA-DR and CK across the subjects could be approximated by Beta distribution respectively with parameters, (1.5, 170) and (1.6, 35) (refer to the Supplementary Material). We used a Gaussian copula (Masarotto and Varin, 2012) to simulate correlated intensity data for two markers which had the above marginal Beta distributions. The simulation strategy was as follows,

1. A random number *I*_*j*_ between 0 and 1 was chosen with probability 0.5 each, respectively standing for group (1), whose subjects had high co-expression of the markers and group (2), whose subjects had mild to none co-expression of the markers. It assigned *j*-th subject to either of the two groups.
2. The intensity vector of two markers, (*X*_1*ij*_, *X*_2*ij*_)^*T*^ for every individual *j* was simulated as follows,
  a. If *I*_*j*_ = 0, simulate a correlation parameter *ρ*_*j*_ from Unif(0.75, 0.9), or else simulate *ρ*_*j*_ from Unif(0, 0.15).
  b. Consider a correlation matrix, 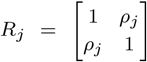 and simulate (*u*_1*ij*_, *u*_2*ij*_)^*T*^ ∼ *N*_2_(0, *R*_*j*_) for *i* = 1, …, *n*_*j*_.
  c. Compute (*v*_1*ij*_, *v*_2*ij*_)^*T*^ = (Φ(*u*_1*ij*_), Φ(*u*_2*ij*_))^*T*^, where Φ() denotes the cumulative distribution function (CDF) of the standard normal distribution.
  d. Perform inverse transformation as, 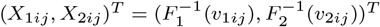, where *F*_1_ and *F*_2_ are Beta distributions with parameters (1.5, 170) and (1.6, 35). Refer to the Supplementary Material for plots of the true joint densities of the markers for two groups of subjects.
3. The clinical outcome of *j*-th subject is simulated as, *Y*_*j*_ = *I*_*j*_*β* + *ϵ*_*j*_, where *ϵ*_*j*_ ∼ *N* (0, *σ*^2^). This step is repeated 100 times to generate 100 different datasets having different *Y* vectors but the same intensity vectors, *X*_1_ and *X*_2_. All the methods are applied on these 100 datasets and empirical power is computed.

Steps 1 − 3 were repeated 20 times and the mean empirical power of different methods were displayed for varying values of the number of cells (*n*_cells_) and the number of individuals (*N*) in Figure 1. The EQMI*\**-based association analysis (EQMI) and the correlation-based association analysis (Corr) achieved comparable performance in all the cases. This particular simulation strategy inherently assumed that the dependence between the markers was linear. Thus, the estimated values of the EQMI*\** and the correlation shared an almost one-to-one relationship making the association analysis using either of them equivalent. Median-Tresholding and Threshold 1 showed similar performance, whereas Threshold 2 had consistently lower power. It showed the importance of choosing a proper threshold in the traditional thresholding-based method for achieving a reasonable performance. All the methods expectedly had the least power when *N* was the smallest, whereas the value of *n*_cells_ did not have any major impact. It implied if the co-expression pattern was well captured even through a smaller number of cells, most of the methods would perform well.

#### 4.1.2 Simulation with squared marker co-expression relationship

The last simulation strategy essentially assumed a linear pattern of co-expression between the markers. Next, we simulated a non-linear pattern of co-expression between the markers where we would expect the correlation-based association analysis (Corr) to perform worse since it could only capture a linear dependence pattern. Steps 1 and 3 of the last simulation strategy were kept the same here and the marker-intensity simulation step i.e., step 2 was changed as follows,

2a. If *I*_*j*_ = 0, simulate *X*_1*ij*_ from Unif(0, 0.1), and *e*_*ij*_ from Unif(0, 0.0005). Construct *X*_2*ij*_ as,

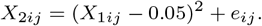
2b. If *I*_*j*_ = 1, independently simulate *x*_1*ij*_, *x*_2*ij*_ both from Unif(0, 0.1).

From Figure 1, we noticed that the EQMI*\**-based association analysis performed the best in all the cases. Threshold 1 and 2 achieved comparable performance for large value of *n*_cells_, whereas Median-Thresholding mostly yielded poor performance. Note that for both the groups of subjects, the correlation between the markers were close to zero as the dependence pattern was non-linear, squared to be specific. Expectedly, the correlation-based association analysis (Corr) had almost no power in every case. The simulation strategy demonstrated why using a generalized measure of co-expression such as EQMI*\** would be more optimal in many cases.

#### 4.1.3 Simulation with circular marker co-expression relationship

In the last simulation setup, the thresholding-based methods performed well despite the marker co-expression pattern being non-linear. Next, we looked into a slightly more complicated co-expression relationship for which the thresholding-based approach would suffer. Steps 1 and 3 of the last two simulation strategies were kept the same here and the step 2 was changed as follows,

2a. If *I*_*j*_ = 0, simulate *X*_1*ij*_ from Unif(0, 0.1), *e*_*ij*_ from Unif(0, 0.0005) and a random number *s*_*ij*_ between -1 and 1. Construct *X*_2*ij*_ as,

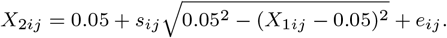
2b. If *I*_*j*_ = 1, independently simulate *X*_1*ij*_, *X*_2*ij*_ both from Unif(0, 0.1).

From Figure 1, we noticed that the EQMI*\**-based association analysis performed the best in all the cases. The correlation-based association analysis (Corr) expectedly performed the worst. Threshold 1, unlike the last simulation, performed significantly worse. In this simulation setup, the subjects of one group had a circular pattern of marker co-expression and the others had almost zero co-expression. Recall that in a two-marker scenario, the thresholding-based methods depended on computing the proportions of the cells positive for both the markers and of the cells positive for only one of the markers (Section 2.1). The difference between these proportions across the two groups of subjects became negligible under this setup which made it difficult distinguishing between them. This explained the overall poor performance of all the thresholding-based methods.

### 4.2 Simulation with three markers

Next, we considered three markers and simulated varied degree of linear dependence between them using Gaussian copula. We only performed the EQMI*\**-based association analysis and the thresholding-based methods in this case. There were again two groups of subjects respectively with high and low co-expression. The simulation strategy was as follows,

1. A random number *I*_*j*_ between 0 and 1 was chosen with probability 0.5 each, respectively standing for groups (1) and (2).
2. The intensity vector of three markers, (*X*_1*ij*_, *X*_2*ij*_, *X*_3*ij*_)^*T*^ for every individual *j* was simulated as follows,
  i. Consider a correlation matrix, 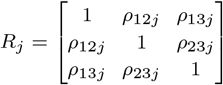. For the subjects in group (1), the off-diagonal elements of the correlation matrix would have high values, whereas they would have low values for the subjects in group (2). We considered three different cases with varying differences between the correlation matrices of the two groups. Case (a) : If *I*_*j*_ = 0, the correlation parameters *ρ*_12*j*_, *ρ*_23*j*_ and *ρ*_13*j*_ were independently simulated from Unif(0.4, 0.6). Otherwise, they were independently simulated from Unif(0.2, 0.4). Case (b) : Regardless of the value of *I*_*j*_, *ρ*_13*j*_ was kept to be 0. If *I*_*j*_ = 0, *ρ*_12*j*_ and *ρ*_23*j*_ were independently simulated from Unif(0.4, 0.6). Otherwise, these two were independently simulated from Unif(0.2, 0.4). Case (c) : Regardless of the value of *I*_*j*_, *ρ*_13*j*_ and *ρ*_23*j*_ were kept to be 0. If *I*_*j*_ = 0, *ρ*_12*j*_ was simulated from Unif(0.4, 0.6). Otherwise, it was simulated from Unif(0.2, 0.4).
  ii. Simulate (*u*_1*ij*_, *u*_2*ij*_, *u*_3*ij*_)^*T*^ ∼ *N*_3_(0, *R*_*j*_) and compute (*v*_1*ij*_, *v*_2*ij*_, *v*_3*ij*_)^*T*^ = (Φ(*u*_1*ij*_), Φ(*u*_2*ij*_), Φ(*u*_3*ij*_))^*T*^.
  iii. Perform inverse transformation as, 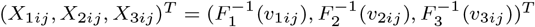, where *F*_1_, *F*_2_ and *F*_3_ are the Beta distributions respectively with parameters (1.5, 170), (1.6, 35) and (1.6, 35).
3. The clinical outcome of *j*-th subject was simulated as, *Y*_*j*_ = *I*_*j*_*β* + *ϵ*_*j*_, where *ϵ*_*j*_ ∼ *N* (0, *σ*^2^). This step was repeated 100 times to generate 100 different datasets having different *Y* vectors but the same intensity data, *X*_1_, *X*_2_ and *X*_3_. All the methods were applied on these 100 datasets and empirical power was computed.

Steps 1 − 3 were repeated 20 times and in Figure 2 the mean empirical power of the methods were displayed. The power of the methods were quite low when *N* was small. The EQMI*\**-based method outperformed both the thresholding-based methods, Threshold 1 and 2 in every case. Threshold 2 had little to no power in most of the cases. Note that, the cases (a), (b), and (c) differed in how different the marker co-expression pattern of the two groups were. The difference between the marker co-expression pattern of the two groups was the largest in case (a) since all the three correlation parameters, *ρ*_12*j*_, *ρ*_23*j*_ and *ρ*_13*j*_ were different across the groups. The difference was the smallest in case (c) as two of the three correlation parameters, *ρ*_23*j*_ and *ρ*_13*j*_ were kept to be 0 in both the groups. Quite expectedly, the power of the methods decreased going from case (a) to case (c), as the difference between the groups of subjects reduced. The decrease was more prominent with Threshold 1 suggesting the method’s lack of robustness.

**Fig. 2.**
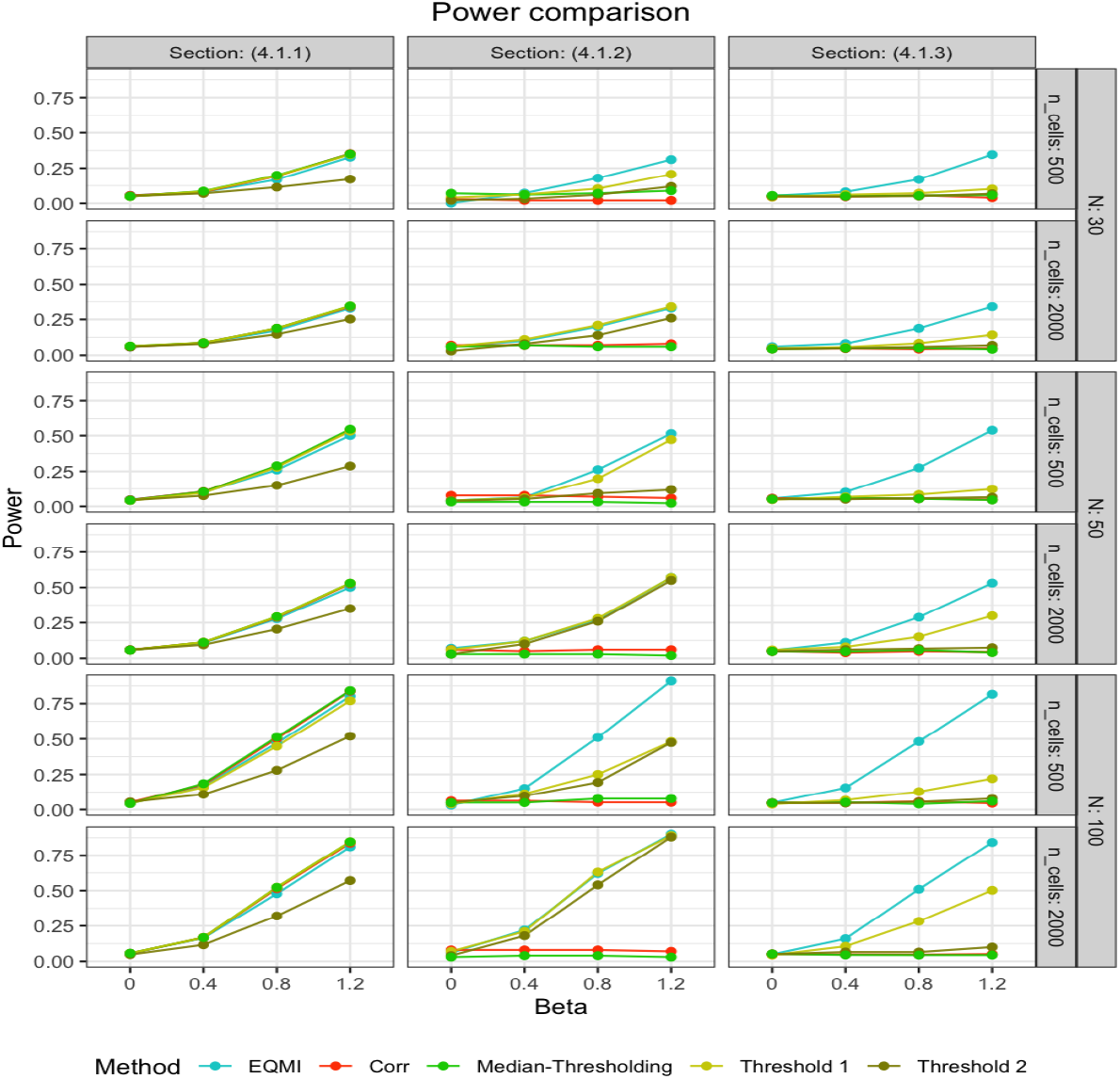
The figure displays the power of different methods under different simulation scenarios from Section 4.1 with two markers for varying numbers of subjects (*N*) and cells (*n*_cells_). On the *x*-axis, the fixed effect size *β* was varied from low to high.

**Fig. 3.**
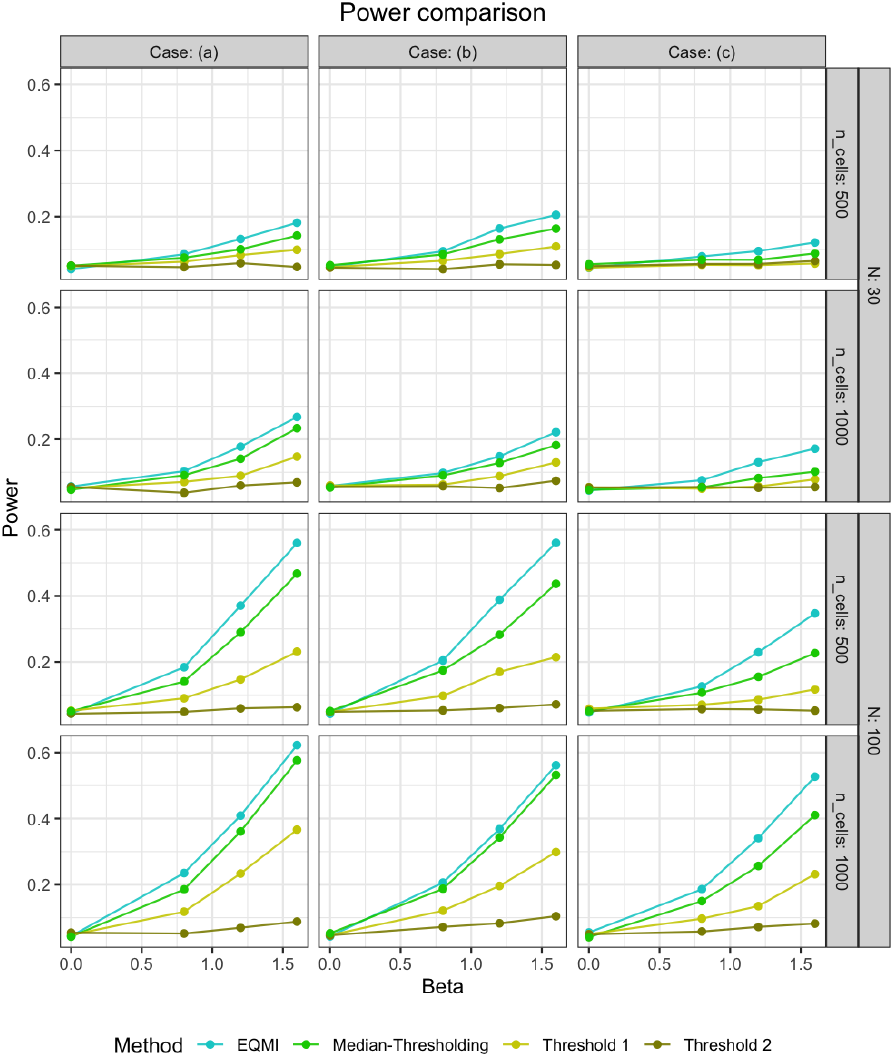
The figure displays the power of different methods under different cases from Section 4.2 with three markers for varying numbers of subjects (*N*) and cells (*n*_cells_). On the *x*-axis, the fixed effect size *β* was varied from low to high.

## 5 Discussion

In multiplex imaging data, studying the interaction or co-expression of multiple functional markers in the cells of the tumor microenvironment (TME) can be crucial for subject-specific assessment of risks. The traditional approach requires a complex step of binarizing the continuous valued marker expression profiles which is prone to subjectivity and can be sub-optimal in many scenarios. The complexity gets exacerbated as the number of markers increases. In this paper, we propose a method for studying co-expression or co-expression of multiple markers based on the theory of mutual information (MI). We treat the subject-specific intensity or expression of every marker as a continuous random variable. We determine how much the markers have co-expressed in the TME of a particular subject by computing a measure known as Euclidean quadratic mutual information (EQMI), comparing the estimated marginal and joint probability density functions (PDFs) of the markers. The formula of EQMI has a similar interpretation as the standard formula of MI but allows a more efficient computation. We adopt and generalize an existing algorithm for computing EQMI that does not require explicitly estimating the joint PDF of the markers, a step which becomes increasingly intractable as the number of markers increases. Next, the subject-level EQMI values are tested for association with the clinical outcomes. The proposed method is free from the subjectivity bias of the traditional thresholding-based method and is readily applicable with any number of markers.

We applied the proposed method to two real datasets, one mIHC lung cancer dataset and one MIBI triple negative breast cancer dataset. In the former, we found high co-expression of the markers, HLA-DR and CK to be associated with the five-year overall survival of the subjects. In the latter, we found high co-expression of the immuno-regulatory proteins, PD1, PD-L1, IDO and Lag3 (IRP’s) to be associated with disease recurrence. We evaluated the performance of our method through several simulation scenarios with two and three markers. In the scenarios with two markers, we showed that all the methods perform well and close to each other if the pattern of dependence (co-expression) between the markers is linear. However, with a more complex non-linear dependence pattern, only the proposed method could achieve respectable power. In the scenarios with three markers, we found that the proposed method performed consistently better than the thresholding-based method and showed superior robustness.

As we have shown in the simulation studies, EQMI can capture both linear and non-linear patterns of co-expression between the markers very well. However, the measure is not well suited for capturing the differences between the patterns. For example, it may happen that one subject has a linear pattern of co-expression, whereas some other subject has a non-linear pattern. The EQMI for both the subjects can be very similar, making it hard to distinguish between them. As a part of our future direction, we would like to improve the method by detecting and incorporating the type of the co-expression pattern. With more than two markers, we studied the co-expression patterns of all possible combinations of the markers and declared significance based on *p*-values corrected by Bonferroni’s method. However, in future, we would like to explore the causal direction between the markers which can then be used to determine a smaller and optimal set of markers and would obviate the need of exploring all possible marker-combinations. In this paper, we have not used any information on the spatial locations of the TME cells. As a future direction, we would like to study the MI between the spatial information and the marker expression profiles with a goal to detect spatially variable markers and their spatial patterns.

Our method is available as an *R* package named MIAMI at this link, https://github.com/sealx017/MIAMI. The package is readily applicable to any multiplex imaging dataset which has cell-level intensity data on two or more markers. In future, we would like to further augment the package’s capability by incorporating a pixel-level analysis as well.

## Acknowledgments

S. Seal acknowledges funding from the Grohne-Stepp Endowment from the University of Colorado Cancer Center, NCI R01 CA129102 and NSF DMS 1914937. D. Ghosh acknowledges funding from NCI R01 CA129102 and NSF DMS 1914937.

