## Supplementary material for "MIAMI: Mutual Information-based Analysis of Multiplex Imaging data"

### 1 Efficient computation of EQMI*

EQMI* for subject $j$ is defined as,

$$\begin{aligned}
&\text{EQMI}^*(X_{1j}, X_{2j}, \ldots, X_{pj}) = \frac{V_J - 2V_C + V_M}{V_J + V_M} \\
&V_J = \int\int \ldots \int f_{12\ldots pj}(x_1, x_2, \ldots, x_p)^2 dx_1 dx_2 \ldots dx_p \\
&V_C = \int\int \ldots \int f_{12\ldots pj}(x_1, x_2, \ldots, x_p) f_{1j}(x_1) f_{2j}(x_2) f_{pj}(x_p) dx_1 dx_2 \ldots dx_p \\
&V_M = \int\int \ldots \int f_{1j}(x_1)^2 f_{2j}(x_2)^2 \ldots f_{pj}(x_p)^2 dx_1 dx_2 \ldots dx_p.
\end{aligned}$$

Estimating the joint density, $f_{12\ldots pj}(x_1, x_2, \ldots, x_p)$ becomes increasingly challenging as $p$ increases. An efficient way of estimating each of the integrals, $V_J, V_C$ and $V_M$ has been discussed in [2] which does not require explicit estimation of the joint or marginal PDFs. However, it has been discussed with a simpler assumption that we generalize further. Before going to the computation, we need to discuss a few key concepts pertaining to multivariate kernel density estimation (KDE). The Parzen KDE of the joint PDF will have the form,

$$\begin{aligned}
&\hat{f}_{12\ldots pj}(\mathbf{x}) = \frac{1}{n_j} \sum_{i=1}^{n_j} w_H(\mathbf{x} - \mathbf{X}_{ij}); \\
&\mathbf{x} = (x_1, x_2, \ldots, x_p)^T, \mathbf{X}_{ij} = (X_{1ij}, X_{2ij}, \ldots, X_{pij})^T;
\end{aligned} \tag{1}$$

where $w_H$ is a multivariate Gaussian kernel with $H$ being the bandwidth matrix. The Parzen KDEs of the marginals will have the forms,

$$\hat{f}_{kj}(x_k) = \frac{1}{n_j} \sum_{i=1}^{n_j} w_{h_k}(x_k - X_{kij}); k = 1, 2, \ldots, p; \tag{2}$$

where $w_{h_k}$ is a univariate kernel with $h_k$ as the bandwidth parameter. Principe (2010) [2] argue that for the joint KDE from Equation 1 to be consistent with the marginal KDEs from Equation 2, the bandwidth matrix $H$ has to be diagonal with elements $h_1, h_2, \ldots, h_p$. It results in a specific structure of the joint KDE,

$$\hat{f}_{12\ldots pj}(\mathbf{x}) = \frac{1}{n_j}\sum_{i=1}^{n_j} w_{h_1}\left(x_1 - X_{1ij}\right) w_{h_2}\left(x_2 - X_{2ij}\right) \ldots w_{h_p}\left(x_p - X_{pij}\right). \tag{3}$$

For $p = 2$, with the form of $\hat{f}_{kj}(x_k)$ from Equation (2) and the form of $\hat{f}_{12\ldots pj}(\mathbf{x})$ from Equation (3) and assuming all the univariate kernels, $w_{h_k}$'s to be Gaussian (denoted by $G$), $V_C$ can be estimated as,

$$\begin{aligned}
\hat{V_C} &= \int\int \hat{f}_{12j}(\mathbf{x})\hat{f}_{1j}(x_1)\hat{f}_{2j}(x_2)dx_1dx_2 \\
&= \int\int \left[\frac{1}{n_j}\sum_{i=1}^{n_j} G_{h_1}\left(x_1 - X_{1ij}\right) G_{h_2}\left(x_2 - X_{2ij}\right)\right] \left[\frac{1}{n_j}\sum_{r=1}^{n_j} G_{h_1}\left(x_1 - X_{1rj}\right)\right] \\
&\qquad\quad \times \left[\frac{1}{n_j}\sum_{s=1}^{n_j} G_{h_2}\left(x_2 - X_{2sj}\right)\right] dx_1dx_2 \\
&= \frac{1}{n_j}\sum_{r=1}^{n_j}\frac{1}{n_j}\sum_{s=1}^{n_j}\frac{1}{n_j}\sum_{i=1}^{n_j}\left[\int G_{h_1}\left(x_1 - X_{1ij}\right) G_{h_1}\left(x_1 - X_{1rj}\right) dx_1\right. \\
&\qquad\quad \left. \times \int G_{h_2}\left(x_1 - X_{1ij}\right) G_{h_2}\left(x_2 - X_{2sj}\right) dx_2\right] \\
&= \frac{1}{n_j}\sum_{i=1}^{n_j}\left[\frac{1}{n_j}\sum_{r=1}^{n_j} G_{\sqrt{2}h_1}(X_{1ij} - X_{1rj})\right]\left[\frac{1}{n_j}\sum_{s=1}^{n_j} G_{\sqrt{2}h_2}(X_{2ij} - X_{2sj})\right].
\end{aligned}$$

For any $p > 2$, the above formula can trivially be generalized as,

$$\begin{aligned}\hat{V}_C &= \int\int\dots\int \hat{f}_{12\dots pj}(\mathbf{x})\hat{f}_{1j}(x_1)\hat{f}_{2j}(x_2)\dots\hat{f}_{pj}(x_p)d\mathbf{x} \\ &= \frac{1}{n_j}\sum_{i=1}^{n_j}\prod_{k=1}^{p}\left[\frac{1}{n_j}\sum_{s=1}^{n_j} G_{\sqrt{2}h_k}(X_{kij} - X_{ksj})\right].\end{aligned} \tag{4}$$

Letting $\hat{V}_k(i,s) = G_{\sqrt{2}h_k}(X_{kij} - X_{ksj})$, $\hat{V}_C$ can be re-written as,

$$\hat{V}_C = \frac{1}{n_j}\sum_{i=1}^{n_j}\prod_{k=1}^{p}\hat{V}_k(i); \hat{V}_k(i) = \frac{1}{n_j}\sum_{s=1}^{n_j}\hat{V}_k(i,s). \tag{5}$$

Similarly $\hat{V}_J$, $\hat{V}_k$ can be estimated as,

$$\begin{aligned}\hat{V}_J &= \frac{1}{n_j^2}\sum_{i=1}^{n_j}\sum_{s=1}^{n_j}\prod_{k=1}^{p}\hat{V}_k(i,s), \\ \hat{V}_M &= \prod_{k=1}^{p}\hat{V}_k; \hat{V}_k = \frac{1}{n_j^2}\sum_{i=1}^{n_j}\sum_{s=1}^{n_j}\hat{V}_k(i,s).\end{aligned} \tag{6}$$

The derivation of the above estimators is discussed in Principe (2010) [2] for a simpler case of $h_k$'s being equal. Equation 5 and 6 enable us to only compute the terms, $\hat{V}_k(i,s)$, for $i, s \in [1,\dots,n_j]$ to directly estimate $\hat{V}_J, \hat{V}_C$ and $\hat{V}_M$ and consequently, $\widehat{\mathrm{EQMI}}^*(X_{1j}, X_{2j},\dots,X_{pj})$. We use Chacón and Duong, 2010 [1]'s diagonal multivariate plug-in bandwidth selection strategy to choose optimal values of $h_k$'s. However, for large $p$ ($p > 6$), Silverman's rule of thumb [3] can be used for choosing $h_k$'s individually to avoid computational deadlock.

#### 2 Additional real data analysis

Table S1 corresponds to the lung cancer data analysis. Tables S2 and S3 correspond to the two sets of markers in the TNBC data analysis.

Table S 1: Comparison of estimated coefficient, hazard ratio (HR), its confidence interval (CI) and LRT $p$-values of the different methods in the association test of five-year overall survival in the mIHC lung cancer dataset.

| Method | Coefficient | HR | CI | $p$-vaue |
|---|---|---|---|---|
| EQMI | -7.264 | 0.0007 | (4.0e-07, 1.238) | 0.0286 |
| Corr | -1.382 | 0.2512 | (0.03131, 2.016) | 0.1838 |
| Median-Thresholding | -0.480 | 0.6189 | (0.3299, 1.161) | 0.1350 |
| Threshold 1 | -0.583 | 0.5585 | (0.3062, 1.019) | 0.0575 |
| Threshold 2 | -1.034 | 0.3555 | (0.1745, 0.7243) | 0.0098 |

Table S 2: Comparison of coefficient estimate and LRT $p$-value of the thresholding-based methods with the markers from set (a) in the MIBI TNBC dataset.

| Outcome | Method | Coefficient | $p$-vaue |
|---|---|---|---|
| Recurrence | Median-Thresholding | 0.596 | 0.152 |
| | Threshold 1 | 0.848 | 0.549 |
| | Threshold 2 | -0.901 | 0.205 |
| Survival | Median-Thresholding | 0.763 | 0.245 |
| | Threshold 1 | 0.337 | 0.106 |
| | Threshold 2 | -0.767 | 0.127 |

Table S 3: Comparison of coefficient estimate and LRT $p$-value of the thresholding-based methods with the markers from set (b) in the MIBI TNBC dataset.

| Outcome | Method | Coefficient | $p$-va |
|---|---|---|---|
| Recurrence | Median-Thresholding | -1.835 | 0.022 |
| | Threshold 1 | -0.710 | 0.255 |
| | Threshold 2 | -0.818 | 0.143 |
| Survival | Median-Thresholding | 0.762 | 0.273 |
| | Threshold 1 | -0.432 | 0.494 |
| | Threshold 2 | -1.025 | 0.090 |

### 3 Simulation density plots

Here, we show the plots of the bivariate joint density of the markers in the subjects from two groups from Section (4.1) of the main manuscript. Note that in every case the subjects from group (1) have high marker co-expression, whereas the subjects from group (2) have little to none co-expression.

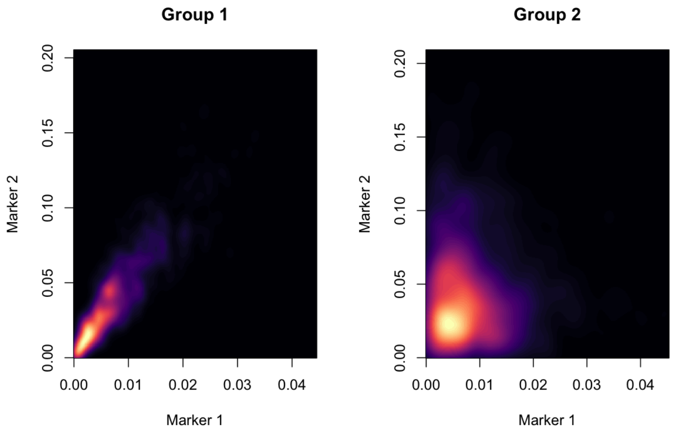


Figure S 1: The right and left panels respectively show the estimated joint density of the markers in a subject from group (1) and a subject from group (2) from Section (4.1.1). A lighter color indicates high value of the estimated density.

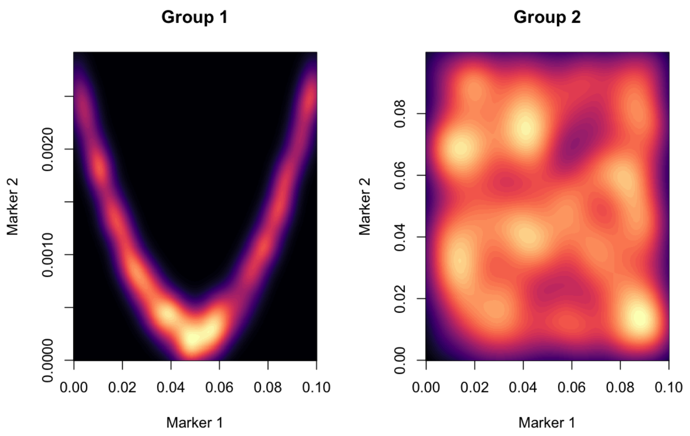


Figure S 2: The right and left panels respectively show the estimated joint density of the markers in a subject from group (1) and a subject from group (2) from Section (4.1.2).

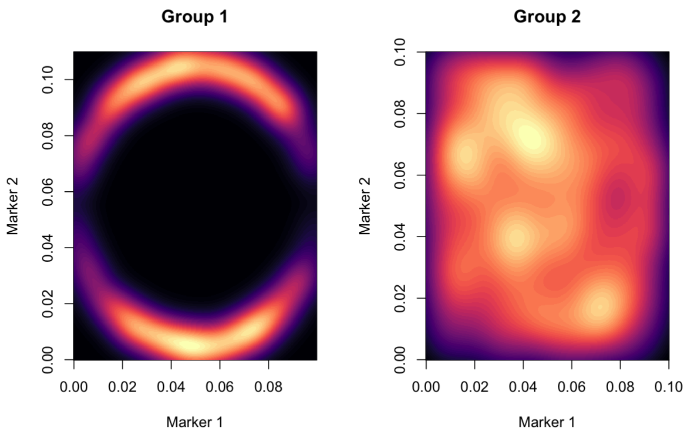


Figure S 3: The right and left panels respectively show the estimated joint density of the markers in a subject from group (1) and a subject from group (2) from Section (4.1.3).
